# Megascale recombination generates millions of catalytically competent PETases

**DOI:** 10.64898/2026.09.18.752806

**Authors:** Pete Heinzelman, Hunter Nisonoff, Akosua Busia, Jennifer Listgarten, Philip A. Romero

## Abstract

Engineering proteins at the scale of modern high-throughput screening technologies remains challenging because most experimental libraries sample only limited regions of sequence space while maintaining relatively low levels of functional diversity. Although recombination-based strategies can introduce substantially larger sequence changes than conventional mutagenesis approaches, it is unclear whether highly recombined enzymes can retain catalytic competence when library sizes approach millions to billions of variants. To investigate this question, we constructed a recombination library by combining ten sequence blocks from ten bacterial PETases, generating a theoretical diversity of 10^10^ chimeric enzymes. Screening of yeast-displayed libraries using a fluorophosphonate probe for catalytic-serine reactivity identified substantial populations of catalytically competent variants despite extensive sequence recombination and dozens of amino acid substitutions per chimera. Approximately 2% of sampled variants possessed active-site serine reactivity, corresponding to an estimated 10^8^ catalytically competent enzymes within the full theoretical library. Deep sequencing showed that functional populations remained broadly distributed across sequence space following selection and contained multiple enriched recombination patterns spanning the PETase structure. Secondary lysate screening and enzyme-normalized activity measurements further identified multiple purified PETase chimeras with soluble esterase activity comparable to or greater than the wild type *Ideonella sakaiensis* PETase on a chromogenic surrogate substrate. Together, these results demonstrate that large-scale recombination can generate millions of diverse and catalytically competent enzymes while remaining compatible with modern high-throughput screening workflows. More broadly, this work establishes recombination-based diversity generation as a scalable strategy for protein engineering and large-scale sequence-function data generation.

## Introduction

Protein engineering is increasingly becoming a data-centric discipline. Advances in machine learning, high-throughput screening, and laboratory automation have created opportunities to learn predictive sequence-function relationships directly from large experimental datasets rather than relying exclusively on low-throughput iterative optimization. At the same time, technologies such as fluorescence-activated cell sorting (FACS), droplet microfluidics [1,2], and magnetic particle-based screening [3] can now evaluate millions to billions of protein variants in a single experiment. Realizing the full potential of these approaches however, requires methods for generating comparably large and structurally meaningful protein diversity while retaining sufficient fractions of functional sequences.

Despite dramatic increases in screening throughput, most protein engineering libraries remain comparatively small and typically explore only narrow local regions of sequence space. This mismatch between library design and screening capacity limits both discovery and large-scale data generation. In particular, while recombination-based approaches can generate substantially larger sequence changes than point mutagenesis and still preserve structural compatibility, previous recombination libraries [4] have generally remained far below the scale accessible to modern screening technologies. As a result, it remains unclear whether recombination can be effectively scaled into the million-to-billion variant regime while maintaining substantial numbers of catalytically competent enzymes.

Here, we address this challenge by constructing and screening a megascale recombination library composed of structurally compatible fragments from ten bacterial PETases [5], yielding a theoretical diversity of 10^10^ chimeric sequences. Using yeast surface display coupled with a serine hydrolase-reactive fluorophosphonate probe [6] and FACS, we found that approximately 2% of sampled variants possessed active-site serine reactivity despite containing dozens of amino acid substitutions relative to their parental enzymes. Secondary screening of enriched clones identified multiple chimeric PETases that could be heterologously expressed in *E. coli* and hydrolyzed a soluble chromogenic PET surrogate substrate at levels comparable to or greater than the *Ideonella sakaiensis* PETase [7]. Together, these results demonstrate that large-scale recombination can generate millions of diverse and catalytically competent enzymes at scales matched to modern high-throughput screening technologies, providing a foundation for scalable protein engineering and large-scale sequence-function data generation.

## Results

### Design and construction of a megascale PETase recombination library

To generate a large and structurally diverse enzyme library compatible with modern high-throughput screening methods, we selected ten bacterial PETases spanning a broad range of sequence diversity while retaining conservation of the canonical α/β-hydrolase fold and catalytic triad architecture. Sequence similarity network analysis showed that the selected enzymes sample diverse regions of PETase sequence space while remaining sufficiently related to support structure-aware recombination (Fig. 1A). Pairwise sequence identities among the selected parents ranged from approximately 56-78%, a regime previously shown to balance sequence diversity and structural compatibility during recombination [4,8]. Together, these enzymes provided a foundation for constructing highly diverse chimeric proteins containing large numbers of amino acid substitutions relative to naturally occurring PETases.

**Figure 1.**
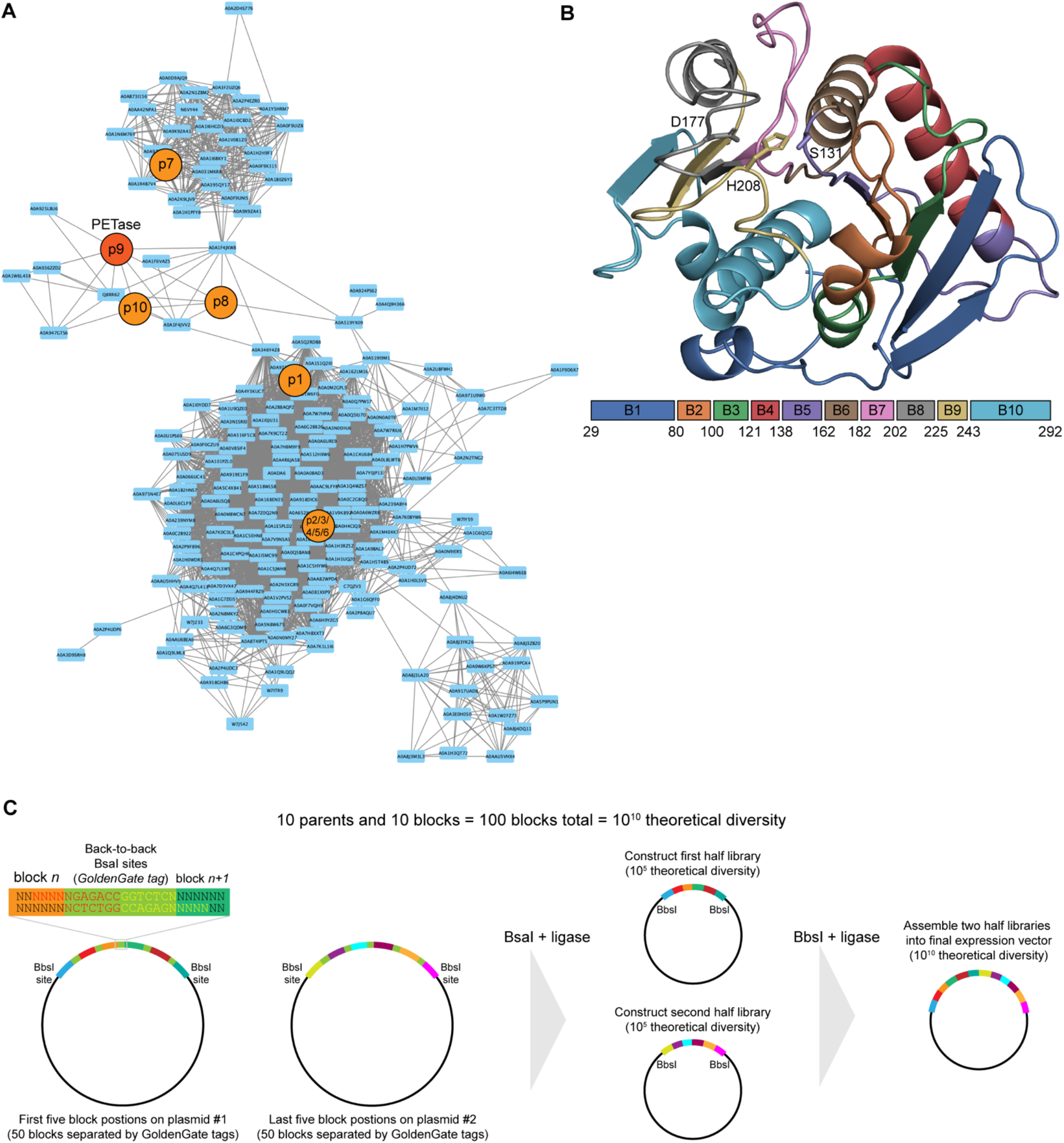
Megascale recombination strategy for generating diverse PETase chimeras. (A) Sequence similarity network of bacterial PETase homologs used for recombination library design. Nodes corresponding to the ten selected parental PETases are highlighted and span broad regions of PETase sequence space while retaining conservation of the canonical α/β-hydrolase fold and catalytic triad architecture. (B) Structure-guided recombination design based on the crystal structure of the *Ideonella sakaiensis* PETase catalytic domain (PDB: 6EQD). The enzyme structure is colored according to the ten contiguous recombination blocks used for library construction. Catalytic triad residues are shown as sticks. Block boundaries were selected to minimize disruption of intramolecular residue contacts while maximizing sequence diversity among resulting chimeras. (C) Modular Golden Gate assembly strategy used to construct the PETase recombination library. Parent-derived recombination blocks were assembled into a yeast surface display vector to generate a library with a theoretical diversity of 10^10^ chimeric PETase sequences.

To design the recombination library, we partitioned the PETase catalytic domain into ten contiguous sequence blocks using structure-aware recombination analysis based on the *Ideonella sakaiensis* PETase structure (PDB: 6EQD). Block boundaries were selected to minimize disruption of intramolecular residue contacts while maximizing sequence diversity among resulting chimeras. The resulting architecture distributed crossover points throughout the enzyme structure, including regions proximal and distal to the catalytic site (Fig. 1B). Recombination of ten blocks from ten parent enzymes yielded a theoretical library diversity of 10^10^ unique chimeric sequences, with individual variants differing by dozens of amino acid substitutions from their closest parental enzymes.

We next developed a modular assembly strategy to construct the recombination library at scales compatible with high-throughput screening (Fig. 1C). Parent-derived DNA fragments corresponding to each recombination block were assembled using Golden Gate cloning and ligated into a yeast surface display vector backbone [9]. Transformation and sequencing analyses indicated successful generation of a large, high-quality library containing approximately 10^5^ unique clones. Both Sanger sequencing of individual variants and Oxford Nanopore sequencing of pooled library DNA showed that >99% of analyzed constructs were full-length chimeras lacking insertions, deletions, or residual restriction enzyme sequences. Representation of individual recombination blocks was broadly uniform across the library, with depletion observed primarily for blocks containing internal restriction sites that interfered with intermediate cloning steps. Together, these results established a structurally diverse and experimentally tractable PETase recombination library suitable for large-scale functional screening.

### High-throughput screening reveals widespread catalytic competence among PETase chimeras

To evaluate whether highly recombined PETase chimeras possessed catalytic competence, we developed a yeast surface display screening assay (Supplementary Fig 1) based on covalent labeling of active-site serines using a fluorophosphonate (FP) probe (Supplementary Fig 2). Yeast-displayed PETases were simultaneously labeled for surface expression and FP reactivity, enabling parallel measurement of protein display and active-site competence by flow cytometry. The wild type *Ideonella sakaiensis* PETase exhibited strong FP labeling relative to a display-positive negative control protein lacking serine hydrolase activity, confirming that the assay could sensitively distinguish catalytically competent PETases from inactive displayed chimeras (Fig. 2A). Because FP labeling reports on the presence of a reactive catalytic serine within a folded active-site environment, this assay enabled high-throughput screening of PETase function at scales inaccessible to conventional substrate-based activity assays.

**Figure 2.**
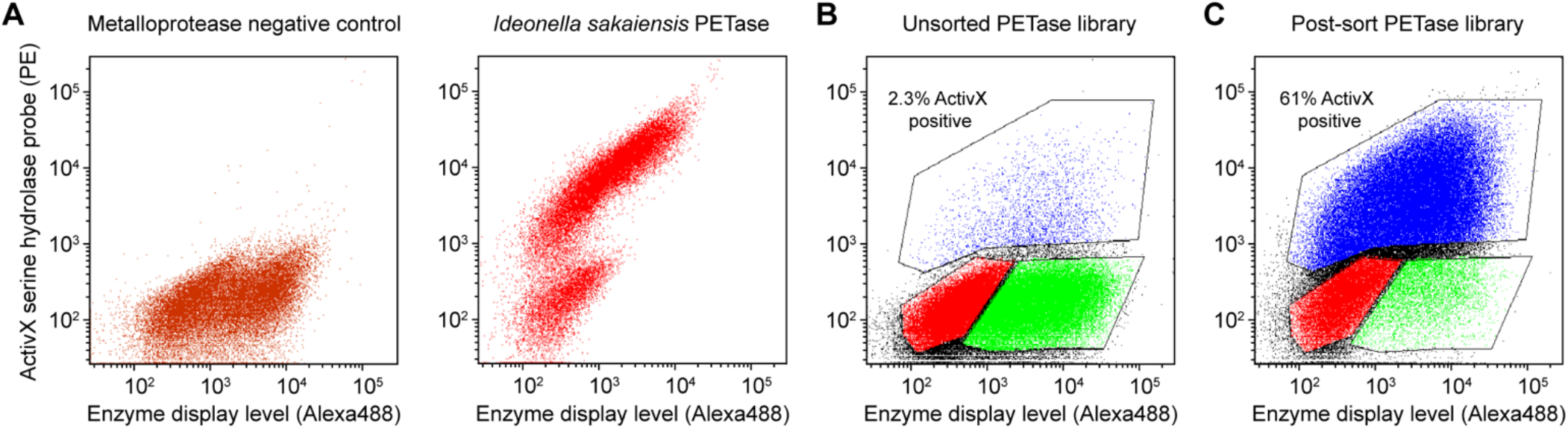
High-throughput screening identifies catalytically competent PETase chimeras. (A) Flow cytometry analysis of yeast-displayed wild type *Ideonella sakaiensis* PETase and a display-positive negative control protein following labeling with fluorophosphonate (FP) probe and streptavidin-phycoerythrin. The wild type PETase exhibited strong FP labeling consistent with catalytic-serine reactivity. (B) Flow cytometry analysis of the unsorted PETase recombination library. Display-positive, FP-reactive yeast corresponding to catalytically competent PETase chimeras comprised approximately 2% of the analyzed population. (C) Flow cytometry analysis of the PETase library following a single round of fluorescence-activated cell sorting (FACS) for display-positive, FP-reactive variants. Sorting enriched the catalytically competent population from approximately 2% to greater than 60% of the total yeast population.

We next applied this assay to the full recombination library. Flow cytometric analysis of the unsorted library revealed a distinct population of display-positive, FP-reactive yeast corresponding to approximately 2% of analyzed variants (Fig. 2B). Given the theoretical diversity of the library, this frequency suggests that the full recombination space may contain on the order of 10^8^ variants retaining catalytic-serine reactivity. Notably, these catalytically competent variants emerged despite individual chimeras containing dozens of amino acid substitutions relative to their closest parental PETases. Although FP labeling does not directly measure PET hydrolysis activity, these results indicate that active-site competence remains remarkably common following large-scale recombination of structurally compatible PETase fragments.

To enrich catalytically competent variants for downstream characterization, we isolated the display-positive, FP-reactive population by FACS. Following a single round of sorting and regrowth, the fraction of yeast exhibiting FP labeling increased from approximately 2% to greater than 60% (Fig. 2C). Because maximal yeast display efficiencies rarely approach 100% [10], additional rounds of sorting were not pursued. Together, these results demonstrate that high-throughput screening can efficiently isolate catalytically competent PETase chimeras from extremely large recombination libraries and that substantial fractions of highly diverse recombined enzymes possess active-site functionality.

### Sequencing analysis identifies recombination patterns associated with catalytic competence

To characterize the composition of catalytically competent PETase populations generated by large-scale recombination, we performed Oxford Nanopore sequencing on both the unsorted starting library and the display-positive, FP-reactive population isolated by FACS. Sequencing analysis showed that the library was composed predominantly of full-length chimeric PETase genes with minimal insertions or deletions, consistent with the high assembly fidelity observed during initial library validation.

Comparison of the unsorted and FACS-enriched populations revealed substantial shifts in recombination block frequencies following screening for catalytic competence (Fig 3AB). Multiple blocks were strongly enriched within the FP-reactive population, whereas others were depleted, indicating that catalytic-serine reactivity depends nonrandomly on recombination composition. Enrichment patterns were distributed across the PETase structure rather than localized to a single region, suggesting that compatibility between multiple structurally distributed sequence elements contributes to the presence of catalytic competence following recombination. Notably, no single parent enzyme dominated the enriched population, consistent with catalytic competence emerging from many distinct combinations of parental sequence fragments.

**Figure 3.**
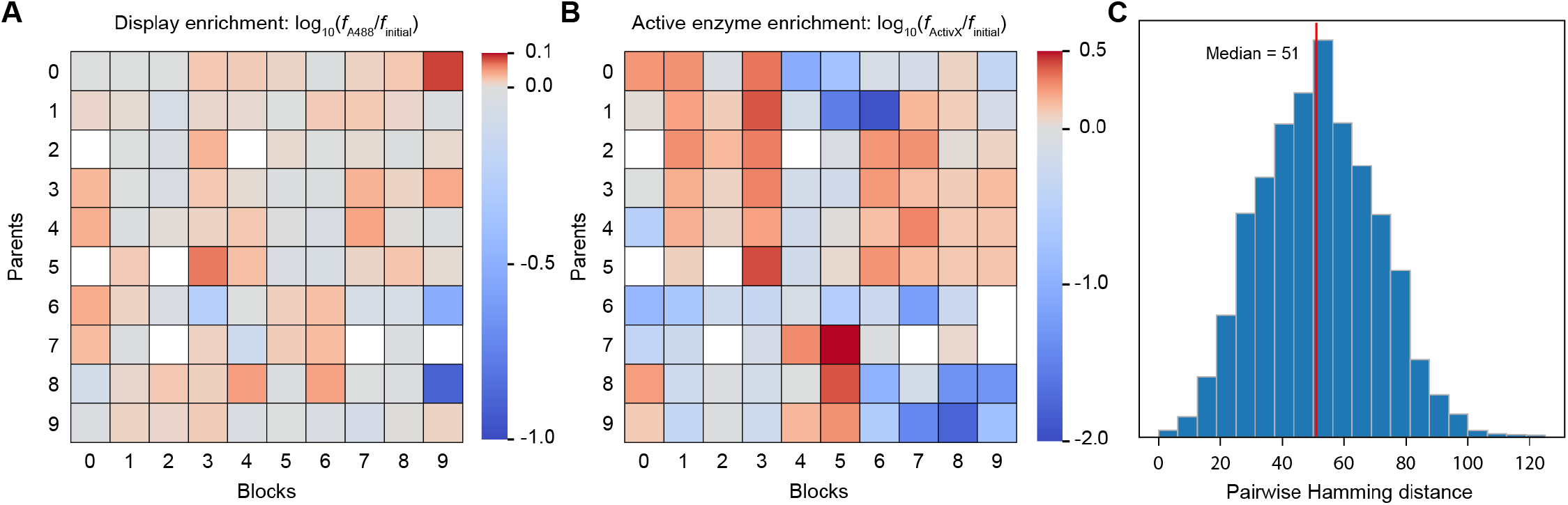
Display and functional enzyme enrichment across the PETase recombination landscape. **(A)** Display enrichment of individual recombination blocks following fluorescence-activated cell sorting (FACS) for surface-displayed (regardless of ActivX active-site probe reactivity) PETase variants. Heat map values represent normalized enrichment of parental recombination blocks in the post-sort population relative to the unsorted starting library. Recombination blocks that were absent from the library due to an inadvertent restriction enzyme site introduced during library construction are shown in white. (**B**) Functional enzyme enrichment of individual recombination blocks following FACS using the ActivX active-site probe. Heat map values represent normalized enrichment of parental recombination blocks in the post-sort population relative to the unsorted starting library. Blocks absent from the library due to the inadvertent restriction enzyme site inclusion in synthesized gene blocks are shown in white. (**C**) The sequence diversity of the sorted FP-reactive sequences. Histogram was generated by sampling random pairs of FP-reactive sequences and calculating their aligned Hamming distance.

Despite strong functional enrichment during FACS, the catalytically competent population remained highly diverse at the sequence level with a median pairwise Hamming distance of 51 (Fig 3C). These findings indicate that large-scale recombination does not converge onto a small number of narrowly optimized solutions but instead generates a broad and heterogeneous population of catalytically competent enzymes. Together, the sequencing analyses demonstrate that high-throughput functional screening can efficiently enrich diverse active PETase chimeras from extremely large recombination libraries while preserving substantial sequence diversity within the selected population.

### Diverse PETase chimeras retain soluble esterase activity after purification

To determine whether catalytically competent PETase chimeras identified by FP labeling possessed activity following soluble expression, we transferred the PETase chimera genes from the FACS-enriched recombination library into an *E. coli* expression system and screened clarified lysates for hydrolysis of the soluble chromogenic ester substrate *p*-nitrophenyl butyrate (pNP-butyrate). This substrate provides a scalable surrogate assay for PETase esterase activity [11] and enables rapid evaluation of large numbers of recombined enzymes following heterologous expression. A total of fifty enriched PETase variants were screened alongside the wild type *Ideonella sakaiensis* PETase and a negative control protein lacking esterase activity [12].

Lysate screening revealed that multiple recombined PETases retained substantial hydrolytic activity toward pNP-butyrate following soluble expression (Fig. 4A). Several variants exhibited activity levels comparable to or greater than the wild type *I. sakaiensis* PETase despite containing extensive sequence recombination across the catalytic domain. Sequence analysis of active variants showed that these enzymes represented multiple distinct recombination patterns rather than convergence onto a single dominant sequence architecture. Together, these results demonstrate that catalytic competence observed during FP-based screening corresponds to recovery of bona fide soluble esterases rather than display-specific artifacts.

**Figure 4.**
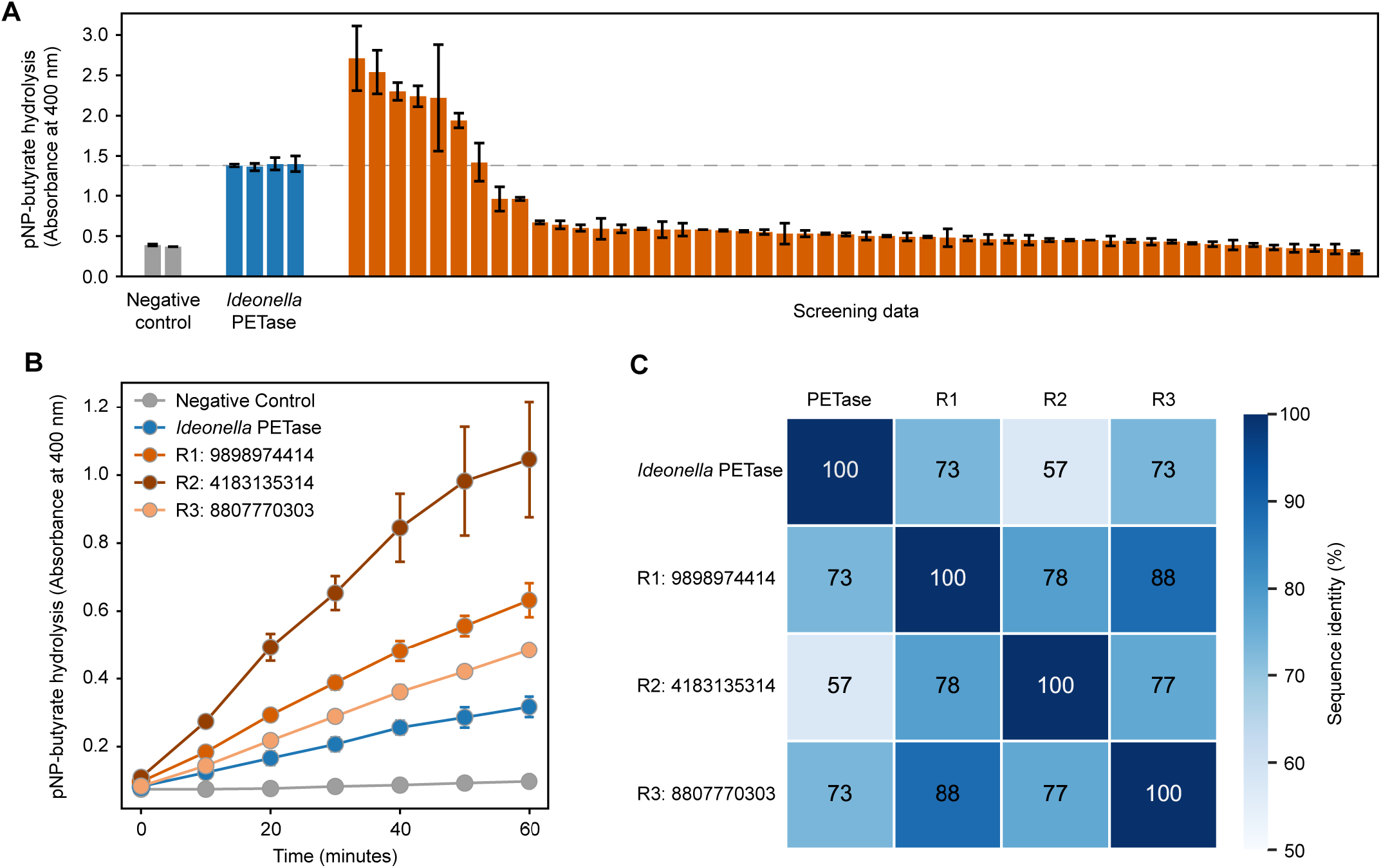
Diverse PETase chimeras retain soluble esterase activity after purification. (A) Lysate-based screening of enriched PETase chimeras for hydrolysis of the chromogenic esterase substrate *p*-nitrophenyl butyrate (pNP-butyrate) following soluble expression in *Escherichia coli*. Activity values are shown relative to the wild type *Ideonella sakaiensis* PETase and a negative control protein lacking esterase activity. Multiple recombined PETase variants retained substantial hydrolytic activity following soluble expression. (B) Enzyme-normalized pNP-butyrate hydrolysis activity measurements for purified PETase-R1 through PETase-R3 and the wild type *I. sakaiensis* PETase. Assays were performed using matched enzyme concentrations following Ni-NTA purification. Several recombined PETase chimeras possessed esterase activity comparable to or greater than the wild type enzyme. (C) Percent sequence identity matrix for wild type *Ideonella* PETase and PETase chimeras R1-R3.

To more rigorously evaluate enzymatic activity, we selected three top-performing variants for purification and enzyme-normalized activity measurements. These purified chimeras, designated PETase-R1 through PETase-R3, were expressed in *E. coli*, purified by Ni-NTA affinity chromatography (Supplementary Fig 3), and assayed at matched enzyme concentrations using pNP-butyrate as substrate. Several purified chimeras retained hydrolytic activity comparable to or greater than the wild type PETase under enzyme-normalized conditions (Fig. 4B). Notably, PETase-R2 and PETase-R3 reproducibly exhibited among the highest activities in both lysate and purified enzyme assays, indicating that functional activity was maintained following purification and concentration normalization.

Although some enriched variants lost activity following purification, potentially reflecting reduced stability or soluble expression efficiency after recombination, multiple highly recombined PETases remained soluble, purifiable, and catalytically active. These findings provide direct evidence that large-scale recombination can generate diverse PETase enzymes that retain functional activity beyond display-based screening assays and remain experimentally tractable following heterologous expression and purification.

## Discussion

In this work, we demonstrate that large-scale recombination of structurally compatible enzyme fragments can generate extensive populations of catalytically competent PETase chimeras at scales matched to modern high-throughput screening technologies. By recombining ten bacterial PETases across ten sequence blocks we constructed a library with a theoretical diversity of 10^10^ variants and found that approximately 2% of sampled sequences retained catalytic-serine reactivity following yeast display screening. Secondary activity assays and purified enzyme measurements further showed that multiple highly recombined chimeras retained soluble esterase activity comparable to or greater than the wild type *Ideonella sakaiensis* PETase. Together, these findings indicate that catalytic competence remains remarkably common following large-scale recombination, even among enzymes containing dozens of amino acid substitutions relative to naturally occurring parents.

Historically, protein engineering has largely operated in a low-diversity regime in which relatively small libraries are explored using iterative mutagenesis and screening [13]. Recombination-based approaches have long been recognized as a powerful strategy for generating larger sequence changes while preserving structural compatibility, but previous studies generally remained limited to libraries far smaller than the capacities of modern screening systems. In contrast, FACS, droplet microfluidics, and related technologies can now evaluate millions to billions of variants in a single experiment. Our results suggest that recombination-based diversity generation can now be scaled into this regime while still maintaining substantial fractions of catalytically competent enzymes. Importantly, the enriched PETase populations remained highly diverse following selection, indicating that functional activity was preserved across many distinct recombination solutions rather than collapsing onto a small number of narrowly optimized sequences.

These findings also have implications for increasingly data-centric approaches to protein engineering. Machine learning methods [14] depend on large and functionally diverse sequence-activity datasets, yet many current experimental workflows remain constrained to relatively local regions of sequence space generated through point mutagenesis. Large-scale recombination provides a complementary strategy for generating broad and structured diversity while maintaining experimentally tractable levels of functional activity. Because recombination simultaneously perturbs many positions throughout a protein, the resulting datasets may be particularly valuable for studying long-range epistatic interactions and training predictive models across broader regions of sequence space than are typically sampled by local mutational scans.

An important feature of this approach is that generation of screening-scale recombination libraries required relatively modest experimental infrastructure and materials cost. Construction of the PETase library described here required only synthesized parent gene plasmids containing Type IIS restriction sites and modular Golden Gate assembly, without iterative cloning workflows or custom primer libraries. As DNA synthesis costs continue to decline, this strategy should be readily extensible to many enzyme families and compatible with increasingly automated and high-throughput protein engineering pipelines. Although PETases were used here as a stringent and societally relevant test case, the overall framework should be broadly applicable to other enzyme classes, such as proteases [15] and laccases [16], for which scalable functional screening assays can be established.

Several limitations of the present study should also be noted. FP labeling reports catalytic-serine competence, rather than direct PET hydrolysis activity, and pNP-butyrate serves only as a soluble surrogate substrate for PETase function. In addition, only a very small fraction of the full theoretical recombination space was experimentally sampled. Nevertheless, even sparse sampling recovered large numbers of catalytically competent enzymes and multiple purified chimeras with substantial esterase activity, suggesting that extensive functional diversity exists within the broader recombination landscape. Future studies combining ultrahigh-throughput screening, automated experimentation, and model-guided library exploration may enable more comprehensive interrogation of these large functional sequence spaces.

Together, these results demonstrate that large-scale recombination can generate diverse populations of catalytically competent enzymes at scales matched to modern high-throughput screening technologies. As screening, automation, and machine learning continue to scale, recombination-based diversity generation may provide a powerful route toward increasingly data-rich and highly scalable protein engineering workflows that enable discovery of improved enzymes, and possibly even entirely new enzyme activities, that cannot be identified using existing methods.

## Materials & Methods

### Selection of PETase parents and recombination design

The wild-type *Ideonella sakaiensis* PETase (UniProt A0A0K8P6T7) and nine additional PET hydrolases were selected from the Plastics-Active Enzymes Database (PAZy). The nine additional parents were the experimentally characterized PET hydrolases with the highest sequence identity to *I. sakaiensis* PETase. Parent sequences were aligned, and signal peptides and N-terminal extensions were removed such that the aligned sequences began at the position corresponding to Q28 of *I. sakaiensis* PETase. The aligned sequences were initially divided into ten blocks of approximately equal amino acid length. Because Golden Gate assembly requires common nucleotide overhangs at each recombination junction, block boundaries were subsequently refined to generate junctions containing two conserved amino acids, one on either side of each breakpoint. When suitable conserved junctions were not naturally present, nearby positions were identified using a Potts model trained on the PETase family and mutated to a common amino acid identity across all ten parents. These junction-standardizing substitutions provided six conserved nucleotides surrounding each breakpoint while preserving compatibility with the family sequence constraints.

The resulting amino acid sequences were reverse translated and codon optimized for *Saccharomyces cerevisiae* using the Twist Bioscience online codon optimization tool. Synonymous codons within the six-nucleotide conserved junction regions were selected to generate compatible four-nucleotide Golden Gate overhangs, with overhang combinations chosen using experimentally measured ligation efficiencies [17]. The 100 resulting fragments (10 block positions × 10 parents) were synthesized as two concatenated constructs: one plasmid containing all parent fragments for block positions 1-5 (50 fragments) and a second containing all parent fragments for block positions 6-10 (50 fragments). Adjacent fragments within each construct were separated by back-to-back BsaI recognition sites, enabling release of the individual blocks with the designed four-nucleotide assembly overhangs. This architecture enabled separate assembly of first- and second-half libraries, each with a theoretical diversity of 10^5^ variants, which were subsequently joined by BbsI-based Golden Gate assembly to generate the full 10-block library with a theoretical diversity of 10^10^ variants.

### Construction of the PETase recombination library

The PETase recombination library was constructed through a two-stage Golden Gate assembly strategy. The 100 parent-derived recombination blocks were synthesized by Twist Bioscience as two concatenated constructs in pTwist Kan high copy plasmids. The first construct contained the 50 fragments corresponding to block positions 1-5 (10 parent-derived sequences per position), and the second contained the 50 fragments corresponding to block positions 6-10. Within each construct, adjacent fragments were separated by back-to-back BsaI recognition sites and designed four-nucleotide overhangs.

Each set of 50 fragments was assembled independently by one-pot BsaI Golden Gate cloning using standard reaction conditions. Digestion liberated the individual parent-derived blocks, and the designed overhangs directed their ordered assembly into five-block chimeras while simultaneously recircularizing the products in the pTwist Kan high copy backbone. Each reaction therefore generated a half-library comprising five block positions with 10 possible parental sequences at each position, corresponding to a theoretical diversity of 10^5^ sequences. The two half-libraries were independently transformed into 10G Supreme electrocompetent *E. coli* (Lucigen), and transformants were pooled and expanded in liquid culture. Plasmid DNA isolated from the pooled populations provided the first- and second-half libraries for subsequent assembly.

The two half-libraries were combined to generate full-length PETase chimeras by BbsI Golden Gate assembly into the yeast surface display vector VLRB.2D-aga2 [9]. BbsI digestion released the five-block inserts from each half-library, and complementary overhangs directed assembly of blocks 1-5 and 6-10 in the correct order into the display vector. The resulting ten-block library had a theoretical sequence space of 10^10^ possible chimeras. Assembly reactions were transformed into 10G Supreme electrocompetent *E. coli*, and pooled transformants were cultured overnight in LB supplemented with carbenicillin. Transformation efficiency estimated by dilution plating indicated that approximately 5 × 10^5^ independent transformants were recovered.

Initial library construction resulted in retention of BsaI-derived assembly sequences in a subset of constructs, consistent with incomplete digestion during the first-stage Golden Gate assembly. The pooled library was therefore subjected to an additional BsaI digestion, ligation, and transformation cycle to remove residual restriction sequences prior to yeast transformation. Library integrity was evaluated by Sanger sequencing of randomly selected clones and Oxford Nanopore sequencing of pooled plasmid DNA. More than 99% of analyzed constructs were full-length chimeric PETase genes without detectable insertions, deletions, or residual restriction sequences.

### Yeast display and fluorophosphonate-based functional screening

The PETase recombination library was transformed into *Saccharomyces cerevisiae* strain EBY100 [3] using a commercial yeast transformation kit (Sigma-Aldrich). Transformants were pooled and cultured in SDCAA medium at 30°C with shaking at 250 rpm. Dilution plating indicated that the transformed yeast population contained >5 × 10^5^ unique transformants.

For induction of yeast surface display, cultures were diluted into SGCAA medium at an initial OD600 of 0.5 and incubated overnight at 20°C with shaking. Approximately 10^7^ induced yeast cells were harvested by centrifugation, washed in phosphate-buffered saline (PBS) containing 0.2% bovine serum albumin (BSA), and incubated for two hours at room temperature with 2 µM ActivX Desthiobiotin-FP Serine Hydrolase Probe (Thermo Fisher Scientific). Following washing, yeast were incubated overnight at 4°C with anti-myc IgY antibody (Aves Labs) to detect surface display. Cells were subsequently labeled with streptavidin-phycoerythrin and Alexa488-conjugated goat anti-chicken IgG secondary reagents for one hour at 4°C.

Flow cytometry and FACS were performed using BD Fortessa and BD FACSAria III instruments located at the University of Wisconsin–Madison Flow Cytometry Core Facility. Surface-displayed PETases were identified by anti-*myc* fluorescence and catalytically competent variants were identified by fluorophosphonate probe labeling. The wild type *I. sakaiensis* PETase served as a positive control for catalytic-serine labeling and a non-serine hydrolase protein [12] served as a negative control. Display-positive, fluorophosphonate-reactive yeast were isolated by FACS and regrown for downstream analysis. Because maximal yeast display efficiencies rarely exceeded ∼75%, additional rounds of sorting were not pursued.

### Sequencing analysis of unsorted and enriched PETase populations

Plasmid DNA was isolated from both unsorted and FACS-enriched yeast populations using the Zymo Research Yeast Plasmid Miniprep II kit. Extracted plasmids were transformed into 10G Supreme electrocompetent *E. coli* and amplified overnight in LB supplemented with carbenicillin. PETase gene inserts were excised by restriction digestion, purified by agarose gel extraction, and submitted to the University of Wisconsin-Madison Next Generation Sequencing Core for Oxford Nanopore sequencing.

Sequencing reads were filtered to retain full-length PETase chimeras lacking insertions or deletions. Individual recombination blocks were assigned by alignment to the ten parental PETase sequences using custom analysis scripts implemented in Python. Block frequencies in unsorted and enriched libraries were calculated by counting the number of reads containing each parental block identity at each recombination position. Enrichment values were calculated as the ratio of normalized post-sort to pre-sort block frequencies.

Pairwise Hamming distance analyses were performed on randomly sampled full-length enriched variants using amino acid sequence alignments. Hamming distances were calculated as the number of amino acid substitutions separating each pair of chimeric sequences.

### Soluble expression and lysate activity screening

Catalytically competent PETase chimeras identified by FACS were transferred into a pET28a cytoplasmic expression vector for soluble expression in *E. coli*. PETase genes were cloned upstream of a C-terminal His_6_ tag and transformed into BL21(DE3) cells. The wild type *I. sakaiensis* PETase and a non-esterase fibronectin domain protein [12] were used as positive and negative assay controls, respectively.

Individual transformants were cultured overnight in LB supplemented with kanamycin and subsequently diluted into fresh media for protein expression. Expression was induced with 1 mM IPTG and cultures incubated for 30 hours at 20°C. Cells were harvested by centrifugation and lysed using Lysonase Bioprocessing Reagent (Sigma-Aldrich). Clarified lysates were obtained by centrifugation and used immediately for esterase activity assays.

Lysate activity screening was performed using the chromogenic esterase substrate *p*-nitrophenyl butyrate (pNP-butyrate). Reactions contained clarified lysate and assay buffer supplemented with pNP-butyrate at a final concentration of 5 mM. Hydrolysis reactions were incubated at 37°C and product formation monitored by absorbance at 400 nm using a Tecan Spark plate reader. Fifty enriched PETase variants were screened alongside wild type and negative control proteins. PETase variants exhibiting activity comparable to or greater than the wild type enzyme were sequenced by Sanger sequencing.

### Purification and enzyme-normalized activity assays

Selected PETase variants identified during lysate screening were retransformed into BL21(DE3) *E. coli* for purification and enzyme-normalized activity measurements. Protein expression cultures were induced with IPTG and incubated for 20 hours at 20°C. Cells were harvested and lysed in PBS containing Lysonase reagent.

Clarified lysates were incubated with Ni-NTA resin (Qiagen) for affinity purification of His_6_-tagged PETases. Following washing with PBS and PBS supplemented with imidazole, bound proteins were eluted with PBS containing 250 mM imidazole and buffer exchanged into EPPS buffer (pH 8.0) using desalting columns. Protein purity and concentration were assessed by SDS-PAGE and densitometry relative to bovine serum albumin standards.

Purified PETases were assayed for pNP-butyrate hydrolysis in 96-well plates using matched enzyme concentrations. Assays were conducted at final enzyme concentrations of 2.5 nM and substrate concentrations of 2.5 mM pNP-butyrate in Tris buffer (pH 7.8). Reactions were incubated at 37°C and absorbance at 400 nm monitored over time. Top-performing purified variants were designated PETase-R1 through PETase-R3.

## Authors’ contributions

PH: Conceptualization, data curation, formal analysis, investigation, methodology, writing of original draft and editing. HN and AB: Conceptualization, investigation and methodology. JL: Conceptualization, resources and supervision. PAR: Conceptualization, data curation, formal analysis, funding acquisition, investigation, methodology, supervision, writing of original draft and editing.

## Code

All software and data analysis code can be found at: https://github.com/RomeroLab/megascale-recombination

## References

1. Roychowdhury H, Romero PA. Microfluidic deep mutational scanning of the human executioner caspases reveals differences in structure and regulation. Cell Death Discov. 2022 10;8(1):7. doi: 10.1038/s41420-021-00799-0.

2. Agresti JJ, Antipov E, Abate AR, Ahn K, Rowat AC, Baret JC, Marquez M, Klibanov AM, Griffiths AD, Weitz DA. Ultrahigh-throughput screening in drop-based microfluidics for directed evolution. Proc Natl Acad Sci U S A. 2010 107(9):4004–9. doi: 10.1073/pnas.0910781107.

3. Chao G, Lau WL, Hackel BJ, Sazinsky SL, Lippow SM, Wittrup KD. Isolating and engineering human antibodies using yeast surface display. Nat Protoc. 2006;1(2):755–68. doi: 10.1038/nprot.2006.94.

4. Otey CR, Silberg JJ, Voigt CA, Endelman JB, Bandara G, Arnold FH. Functional evolution and structural conservation in chimeric cytochromes p450: calibrating a structure-guided approach. Chem Biol. 2004 11(3):309–18. doi: 10.1016/j.chembiol.2004.02.018.

5. Lusty Beech J, Clare R, Kincannon WM, Erickson E, McGeehan JE, Beckham GT, DuBois JL. A flexible kinetic assay efficiently sorts prospective biocatalysts for PET plastic subunit hydrolysis. RSC Adv. 2022 12(13):8119–8130. doi: 10.1039/d2ra00612j.

6. Rajagopalan S, Wang C, Yu K, Kuzin AP, Richter F, Lew S, Miklos AE, Matthews ML, Seetharaman J, Su M, Hunt JF, Cravatt BF, Baker D. Design of activated serine-containing catalytic triads with atomic-level accuracy. Nat Chem Biol. 2014 10(5):386–91. doi: 10.1038/nchembio.1498.

7. Austin HP, Allen MD, Donohoe BS, Rorrer NA, Kearns FL, Silveira RL, Pollard BC, Dominick G, Duman R, El Omari K, Mykhaylyk V, Wagner A, Michener WE, Amore A, Skaf MS, Crowley MF, Thorne AW, Johnson CW, Woodcock HL, McGeehan JE, Beckham GT. Characterization and engineering of a plastic-degrading aromatic polyesterase. Proc Natl Acad Sci U S A. 2018 115(19):E4350–E4357. doi: 10.1073/pnas.1718804115.

8. Li Y, Drummond DA, Sawayama AM, Snow CD, Bloom JD, Arnold FH. A diverse family of thermostable cytochrome P450s created by recombination of stabilizing fragments. Nat Biotechnol. 2007 25(9):1051–6. doi: 10.1038/nbt1333.

9. Tasumi S, Velikovsky CA, Xu G, Gai SA, Wittrup KD, Flajnik MF, Mariuzza RA, Pancer Z. High-affinity lamprey VLRA and VLRB monoclonal antibodies. Proc Natl Acad Sci U S A. 2009 106(31):12891–6. doi: 10.1073/pnas.0904443106.

10. Burns ML, Malott TM, Metcalf KJ, Puguh A, Chan JR, Shusta EV. Pro-region engineering for improved yeast display and secretion of brain derived neurotrophic factor. Biotechnol J. 2016 11(3):425–36. doi: 10.1002/biot.201500360.

11. Heyde SAH, Arnling Bååth J, Westh P, Nørholm MHH, Jensen K. Surface display as a functional screening platform for detecting enzymes active on PET. Microb Cell Fact. 2021 20(1):93. doi: 10.1186/s12934-021-01582-7.

12. Heinzelman P, Low A, Simeon R, Wright GA, Chen Z. De Novo Isolation & Affinity Maturation of yeast-displayed virion-binding human fibronectin domains by flow cytometric screening against Virions. J Biol Eng. 2019 Oct 13:76. doi: 10.1186/s13036-019-0203-2.

13. Morawski B, Quan S, Arnold FH. Functional expression and stabilization of horseradish peroxidase by directed evolution in Saccharomyces cerevisiae. Biotechnol Bioeng. 2001 76(2):99–107. doi: 10.1002/bit.1149.

14. Freschlin CR, Fahlberg SA, Romero PA. Machine learning to navigate fitness landscapes for protein engineering. Curr Opin Biotechnol. 2022 Jun;75:102713. doi: 10.1016/j.copbio.2022.102713.

15. Mikolajczyk BM, Golinski AW, Hackel BJ. Enzyme-substrate co-display on yeast empowers engineering of tobacco etch virus protease activity. Protein Eng Des Sel. 2025 38:gzaf011. doi: 10.1093/protein/gzaf011.

16. Lee SY, Roh H, Gonzalez-Perez D, Mackey MR, Hoces D, McLaughlin CN, Lin C, Adams SR, Nguyen K, Kim KY, Luginbuhl DJ, Luo L, Udeshi ND, Carr SA, Hernández-López RA, Ellisman MH, Alcalde M, Ting AY. Directed evolution of LaccID for cell surface proximity labeling and electron microscopy. Nat Chem Biol. 2025 21(12):1895–1905. doi: 10.1038/s41589-025-01973-6.

17. Pryor JM, Potapov V, Kucera RB, Bilotti K, Cantor EJ, Lohman GJS. Enabling one-pot Golden Gate assemblies of unprecedented complexity using data-optimized assembly design. PLoS One. 2020 15(9):e0238592. doi: 10.1371/journal.pone.0238592.

