## Supplementary Information for "Megascale recombination generates millions of catalytically competent PETases"

**Supplementary File**

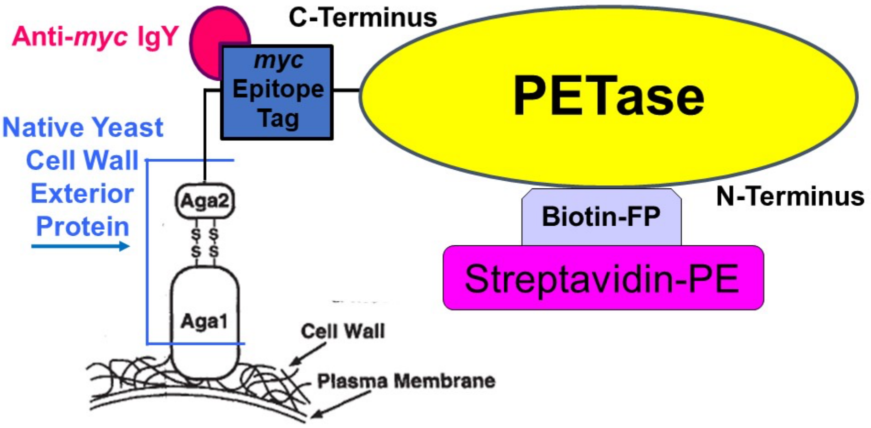


**Supplementary Fig 1.** Yeast display schematic. ACE2 C-terminus linked to Aga2 native yeast surface protein on yeast cell wall. Anti-*myc* IgY (from chicken) used to quantify ACE2 display. Anti-*myc* IgY is not fluorescently labeled; fluorescent detection of *myc* epitope tag achieved via incubation with Alexa488-conjugated anti-chicken IgG (from goat) as secondary label (not depicted). Covalent bond formation between biotin-FP probe and PETase nucleophilic Ser detected with streptavidin-phycoerythrin secondary label. Each yeast cell displays up to $10^4$ copies of a single ACE2 variant on its surface. Schematic adapted from Boder and Witrup. *Nat Biotechnol.* 1997 15(6):553-7.

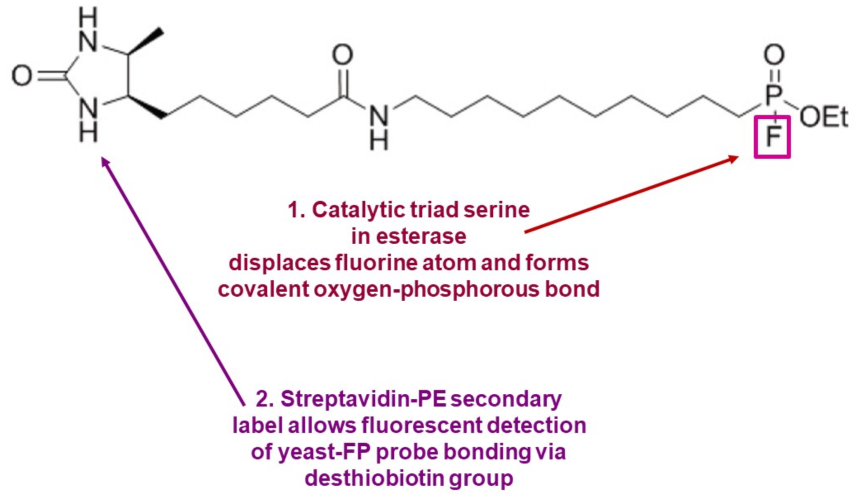


**Supplementary Fig 2.** Biotin-FP serine hydrolase probe (ThermoFisher catalog #88317)

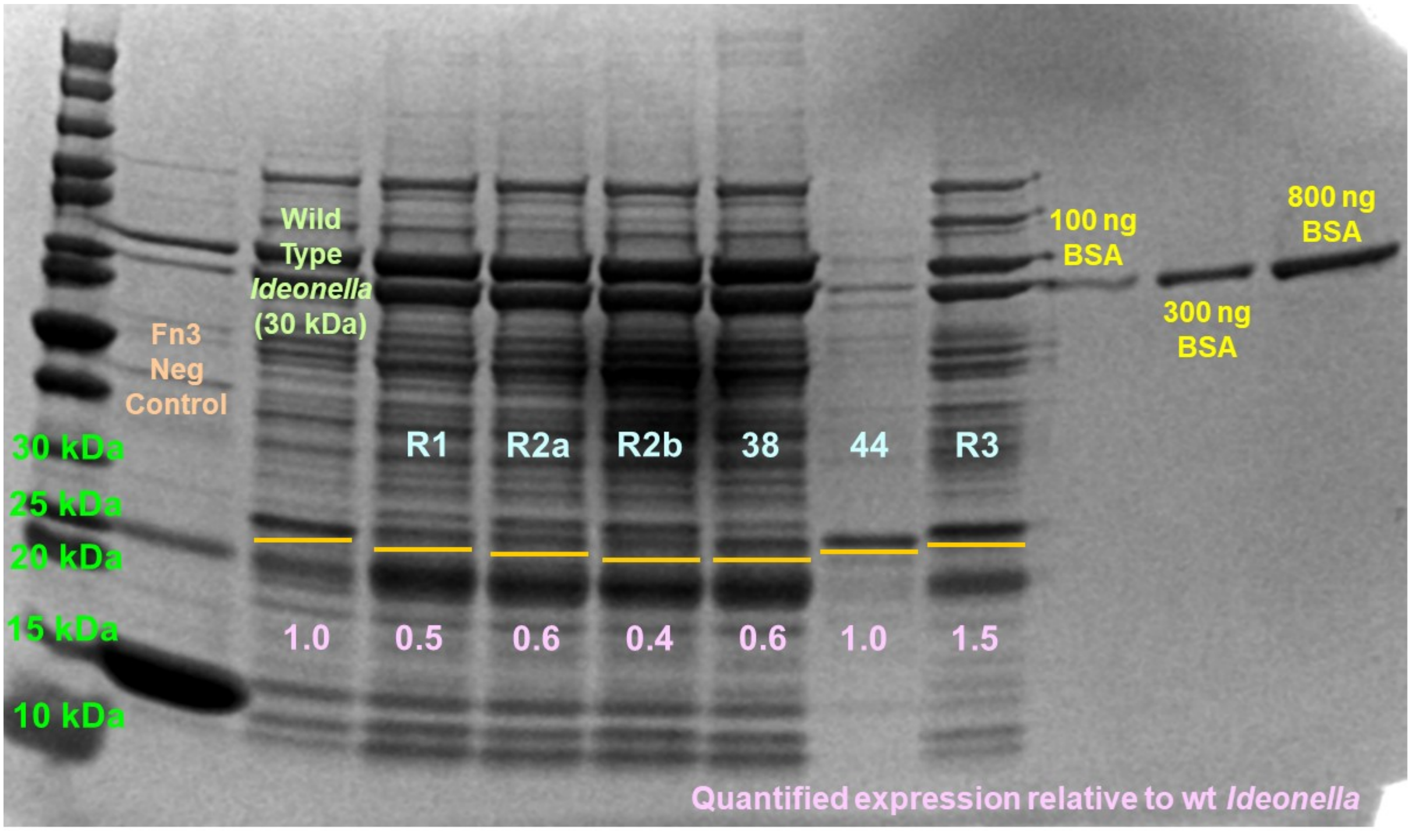


**Supplementary Fig 3.** Post-purification SDS-PAGE analysis of leading PETases identified in pNP-butyrate hydrolysis screen of Fig 4a. *Ideonella* and chimeric PETase anticipated molecular weight is ~ 30 kDa. Numbers marking lanes denote clone numbers for PETase chimeras. Clones R2a and R2b were identified as identical (block sequence 4183135314) by DNA sequencing. R2a protein prep was used in Figure 4b activity assay. Orange lines appear underneath bands that were quantified using GelAnalyzer 23.1.1 to estimate PETase concentration. Pink numbers denote quantified band intensities relative to wild type *Ideonella*. Purified protein for PETase chimeras 38 and 44 (clone numbers for lysate screen of Fig 4a) were not active toward pNP-butyrate.
